# Distribution of Entomopathogenic Nematodes in Some Selected Insect Species in Osogbo Local Government Area of Osun State, Nigeria

**DOI:** 10.1101/2023.02.23.528628

**Authors:** Q.O Adeshina, A.M Rufai, O.A Surakat, S.O Nzeako

**Affiliations:** Parasitology Unit, Department of Zoology, Faculty of Basic and Applied Sciences, College of Science and Engineering, P.M.B. 4494, Osun State University, Osogbo, Nigeria; Parasitology and Public Health Unit, Department of Animal and Environmental Biology, University of Port Harcourt, Nigeria

**Keywords:** Biological control, Distribution, Entomopathogenic nematodes, *Heterorhabditis*, *Mermis* spp., Infectivity, Insect pests and Occurrence

## Abstract

Entomopathogenic nematodes (EPNs) are leading biological control agents used to combat many insect pests in many regions of the world. This study investigated the distribution of entomopathogenic nematodes in four insect species; *Zonocerus variegatus, Mantis religiosa*, Butterfly and Moth across dry and rainy seasons in Osogbo Local Government Area of Osun State. Insect samples were collected from different sampling stations (cultivated farmlands, vegetation of grasses, and forested lands) using an entomological sweep net. The insects were dissected in a normal saline medium for the presence of EPN. A further verification was made, 60 soil samples were retrieved randomly from the disturbed agroecosystem (where insects were sampled) and an undisturbed agroecosystem (Osun-Osogbo Groove). The soil samples were screened for EPN presence using *Tenebrio molitor* larva (mealworm) as baits, while infected baits are placed on modified white traps to recover EPNs. A total of 876 insects samples comprising; *Zonocerus variegatus* 556 (63.5%), *Mantis religiosa* 36 (4.1%), Butterflies 158 (18.0%) and Moths 126 (14.4%) were collected from the disturbed agroecosystem based on sweep net efficiency and species availability. After screening, only 1 (0.1%) insect specimen (*Mantis religiosa*) specimen successfully harbour an EPN, identified as *Mermis* spp. Result from statistical analysis indicates that both season and insects age do not have influence on the distribution of EPN (P>0.05). Moreover, the low infectivity of EPN in the sampled insect was presumed to be caused by EPNs’ foraging strategies, flooding and other host of factors. This led to further probing through screening of soil samples. Of all 36 soil samples screened from the disturbed agroecosystem, EPNs had zero prevalence. However, from all the (24) soil samples collected from the undisturbed agroecosystem, EPNs belonging to the genus *Heterorhabditis* were recovered and identified using morphological and morphometric characters. The absolute frequency of occurrence of EPN in the disturbed agroecosystem was zero compared to 100% recorded for the undisturbed agroecosystem. This study infers that EPN occurrence, dispersal, and persistence in the ecosystem are found to be adversely affected by intense anthropogenic activities.

## 1. INTRODUCTION

Insect pests’ invasion has become a major setback in achieving sustainable crop production and, thus, leading to food insecurity in Nigeria and other sub-Saharan African countries (Ozeum *et al*., 2021). Insect pests cost farmers billions of dollars each year in crop losses and the cost of controlling them (Sharma *et al*., 2017). The use of chemical insecticides has been the leading measure incorporated to combat an array of insect pests (Isman, 2019). Many are non-biodegradable, contaminate the environment, harm humans and other non-target organisms (Ozeum *et al*., 2021). The rising demand for organic food calls for biocontrol alternative to pest control by farmers (Das *et al*., 2020). Biological insecticides are screened as viable alternatives or valuable complements in integrated pest management when used appropriately (Stankovic *et al*., 2020). Research has shown that biological insect control is target specific, eco-friendly, and has a low possibility of pest resistance development (Poinar, 1990; Kaur & Garg, 2014; Kwenti, 2017).

A specific group of nematodes known to parasitize insects are called insect pathogenic or entomopathogenic nematodes or entomophagous nematodes (EPN) (Poinar, 1990). Entomopathogenic nematodes (EPNs) are soil-inhabiting parasites of insect pests. They have been discovered on every continent except Antarctica and in various ecological settings, ranging from cultivated fields to deserts (Abate *et al*., 2017; Bhat *et al*., 2020). EPNs are non-toxic to humans, safe for the environment, can be mass-produced, formulated, and applied easily (Lacey & Georgis, 2012). They have been verified to suppress the activities of insect pests in many agrarian settings, and thereby making EPNs, excellent biological control agents for many economically important insect pests (Al-Zaidawi *et al*., 2020). The Orders: Mermethida, Aphelenchida, Tylenchida, and Rhabditida, have been the most intensively studied of all orders of nematodes associated with insects. However, only the Rhabditid genera; *Heterorhabditis* and *Steinernema* are widely used for insect control due to their high infectivity, pathogenicity, and easy manipulation (Mracek, 2002). EPNs invade natural insect openings such as the mouth, anus, spiracles, and cuticles, travel to the hemocoel, and release symbiotic bacterial cells, which proliferate and kill the insects within 48 hours while simultaneously providing nutrition and optimum conditions for nematode development and reproduction (Bahadur, 2018; Liu *et al*., 2020). These nematodes–bacteria complexes are incredibly lethal to insects and are widely recognized as one of the most effective non-chemical insect pest control strategies. However, members of other families, such as Mermithidae, do not have a symbiotic association with entomopathogenic bacteria (EPB). The Mermithids use their stylet to penetrate the insect cuticle, then migrate to the host hemocoel to initiate their parasitic phase (Devi, 2020).

EPNs suitability for commercialization is often influenced by their host range, which revolves around EPNs’ foraging strategies, ability to penetrate different insect hosts, and natural population density, which significantly affects their occurrence, dispersal, and persistence in the field. The host range of most known EPN species remain unknown, even though most species have been isolated from soil samples using the highly susceptible wax moth; *Galleria mellonella* larvae or mealworm; *Tenebrio molitor* larvae as bait insects (Shapiro-Ilan *et al*., 2018). Although many EPN species have infected many insect species in laboratory assays, the host range is much narrower in the field due to the ecology of the nematodes and their potential hosts. In spite of the successful in vivo determination of EPNs host range, field determination is influenced by ecological factors which are variable.

There are concerns about introducing exotic entomopathogenic nematodes to combat insect pest’s invasion because they may harm non-target organisms (Barbosa-Negrisoli *et al*., 2009). Surveys have been conducted in almost all parts of the world to isolate locally adapted EPN species or isolates to formulate and commercialize them (Hominick, 2002). Also, several surveys for EPNs have been documented with several new species and strains from Africa (Bhat *et al*., 2020). However, studies on the occurrence and distribution of indigenous EPN species in Nigeria and their potential use as biocontrol agents on field trials is still at an early stage (Akyazi *et al*., 2012; Rufai *et al*., 2020). The use of EPNs will boost IPM in Nigeria, as well as proffering solutions in terms of cost and accessibility, especially for peasant farmers in many regions of the country dealing with high cost and difficulty in procuring pesticides, and perhaps increase food production, food security and adding positive impact on the Nigerian economy. However, the potential use of EPNs as a veritable biocontrol agent of insect pests in Nigeria can only be ascertained by understanding the dynamics of EPN occurrence and their suitability to infect target insect pests in the country (Yan *et al*., 2018).

Therefore, the need for a survey on the pathogenicity of EPN local strains on insect pests in the natural ecosystem in Osun State is vital for the successful use of EPN as a safer alternative to chemical insecticides. This survey will form baseline information for the prevalence of EPNs in some insect pests, their level of infectivity, and the suitability of these local stains to target insect pests in Osun State. This present study aims to determine the distribution of local strains of EPNs in some insect species in the Osogbo Local Govt. Area of Osun Sate, Nigeria.

## 2. MATERIAL AND METHODS

### 2.1 Study Area

The study was conducted in the Osogbo Local Government Area of Osun State, Southwestern Nigeria. The Local Government Area serves as an administrative centre for the local government council and the state capital of Osun State. It has a population of 156,694 people with an area of 47km^2^, according to the Population and Housing Commission Census (2006). The LGA lies within coordinates 7.46^0^N and 4.34^0^E. The sampling points include cultivated farmlands, the vegetation of grasses, and forested lands.

### 2.2 Collection of Insect Samples

Insect samples were obtained from the field using traps based on the rate of catch and species accessibility. Variegated grasshopper; *Zonocerus variegatus*, Praying Mantis; *Mantis religiosa*, Butterfly and Moth are the four insect species sampled. Using the field technique described by (Colwell, 2012), an entomological sweep net of 40cm diameter aperture and 80cm depth was deployed to catch flying insect samples. Captured insects were kept in holed plastic containers to allow air passage and transported to the laboratory for further analysis.

### 2.3 Evaluation of Phenotypic Characteristics of Insect Samples

Following (Ozeum *et al*., 2021), the weight (g) of the insect samples was weighed using an electronic measuring scale, while a meter rule was used to measure the standard and total lengths (cm) of the insects. The standard length of the samples involves the length of the insect, excluding the jumping legs, while the total length includes the stretched-out jumping legs of the insects. Following Stubins (2015), morphological distinctions were used to determine the age of the insects. The age of the insects was determined by the development of the wings. The wings of the nymph stage used in the study were either underdeveloped or none existent, while the adult stages have fully developed wings.

### 2.4 Dissection and Parasitological Screening of the Insect Samples

Following Stubins (2015), a drop of 70% ethanol was placed on a dissecting board to euthanize freshly caught insect samples before dissection, whereas previously caught insect samples were refrigerated. The samples were placed on dissecting board with their ventral sides facing upward, and their legs were cut off. Their abdomens were slightly raised before executing an incision in the middle to reveal the abdominal contents. The lateral portion of the abdominal coverings was pulled apart with forceps and pinned to the dissecting board to expose the digestive tract, which was examined in situ using a magnifying lens. For about 20 minutes, the dissected insect was allowed to stand overwhelmed in normal saline to detect any nematode emergence. Then, the abdominal fluid and scrapings of the gut endothelium were placed on a wash glass and examined under a dissecting microscope to identify nematodes. The gut endothelial scrapings and abdominal fluid were afterward used to prepare a saline wet mount to identify nematode or helminths’ eggs, which were stained with lugos iodine thereafter for clarity (Cheesborough, 1987).

### 2.5 Collection of Soil Samples

A total of 60 soil subsamples were collected for screening. All soil samples were collected randomly at a depth of 20cm with a hand trowel and stored in a polythene bag. The samples were labeled accordingly using a waterproof marker with site information. Thirty-six (36) soil subsamples (at four subsamples per sampling station) were collected in a disturbed agroecosystem, where the insect samples were captured. The subsamples were bulked and mixed thoroughly for each sampling station using a hand trowel to form a composite sample. Then, twenty-four (24) soil subsamples (at three subsamples per sampling station) were collected from an undisturbed agroecosystem (Osun-Osogbo Groove), which falls within the Osogbo Local Govt. Area. Also, the subsamples were bulked and mixed thoroughly for each sampling station using a hand trowel to form a composite sample. The soil samples were transported in a cold chain to maintain standard temperature range.

### 2.6 Extraction of EPNs From Soil Samples

*Tenebrio molitor* larvae were initially obtained from the Zoological Garden, Osun State University, Osogbo, Nigeria. Each composite soil sample were placed in a 500 ml plastic container, moistened, baited with three mealworm larvae, covered with lid and incubated in the laboratory at room temperature (Bedding & Akhurst, 1975) (Figure 1; A, B, C & D). The containers were checked after every 24hours for larval mortality (Figure 1E). The cadavers collected were rinsed with distilled water to remove soil particles and then transferred to a modified White trap (Kaya & Stock, 1997) (Figure 1F). Infective juveniles that migrate from the dead mealworm larvae were collected regularly from the white traps. Each nematodes isolate was cultured on fresh mealworm larvae and then rinsed and stored in a Tetrapak containers.

**Figure 1:**
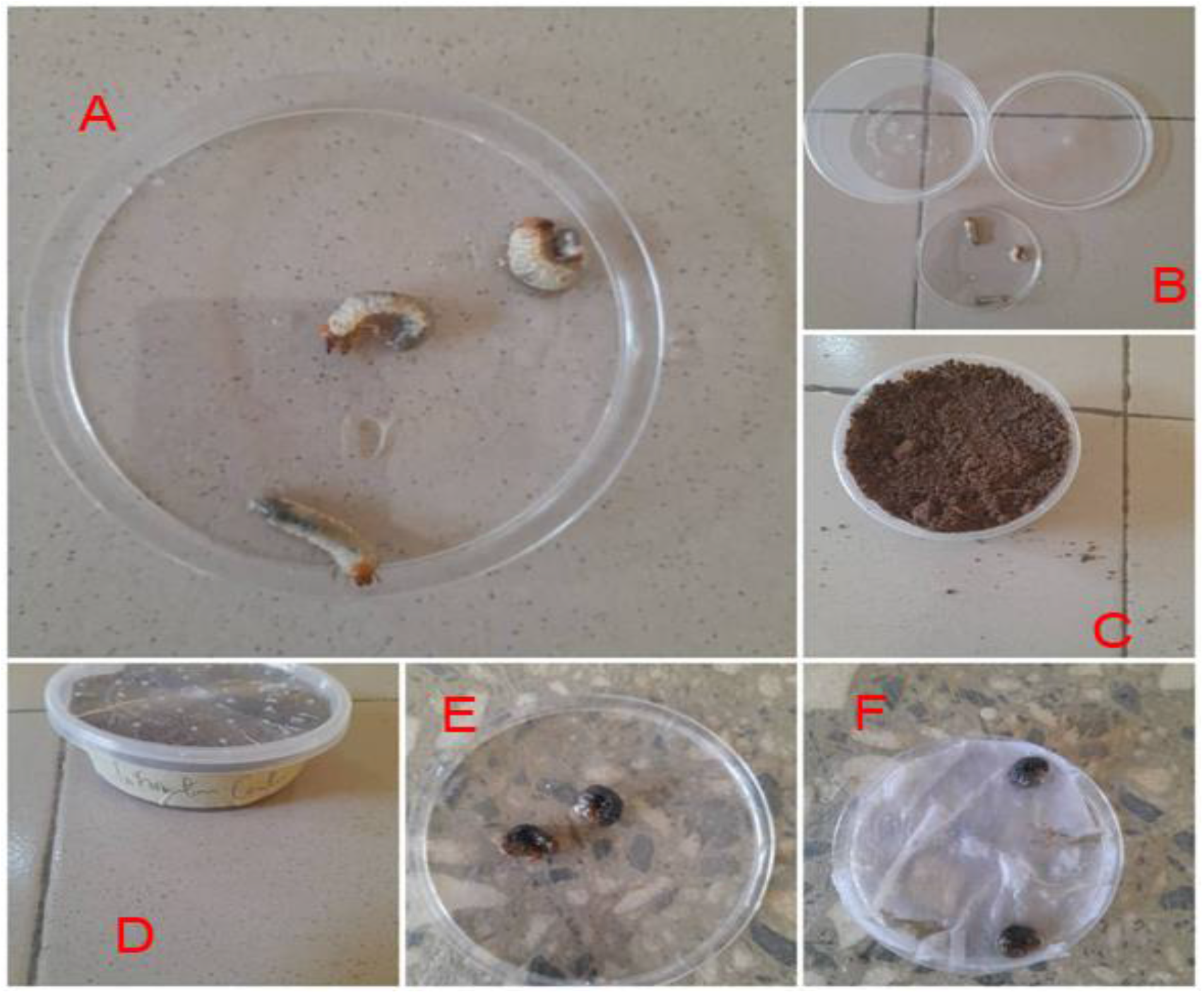
Procedure involving EPN recovery from soil samples.

### 2.7 Identification of EPNs

EPN isolates were fixed and heat-killed using equal volume of triethanolamine–formalin (TAF) fixative (Hazir *et al*., 2022). EPN isolates were picked with a needle and transferred to watch glass with 0.5 mL anhydrous glycerol solution Then, EPN isolates were identified using the morphological and morphometric features comprising; L = length, W = greatest body diameter, EP = distance from anterior end to excretory pore, NR = distance from anterior end to nerve ring, ES = pharynx length, T = tail length with second stage cuticle, D% = EP/ES × 100, E% = EP/T × 100, of the infective juveniles and second generation adults under a digital compound microscope (Baker & Capinera, 1997; Hazir *et al*., 2022).

### 2.8 Frequency of EPNs Occurrence

The absolute frequency of entomopathogenic nematode in soil samples from both disturbed and undisturbed agroecosystems were calculated to assess the EPN’s population. Absolute frequency = No. of samples containing entomopathogenic nematode species /Total number of samples collected × 100

### 2.9 Statistical Analysis

The data obtained were analyzed with the Student t-test with a p-value of ≤ 0.05 and measures of central tendency using Microsoft Excel Software and Statistical Package for Social Sciences (SPSS).

## 3. RESULTS

### 3.1 Prevalence of Entomopathogenic Nematodes (EPNs) in Sampled Insects in the Study

A total of 876 insects were collected for the study from different sampling stations within the territory of the Osogbo Local Govt. Area of Osun State. Out of the 876 insects collected for the survey, 556 (63.5%) were *Z. variegatus* with zero infection, 36 (4.1%) were *M. religiosa* with 2.8% infected, 158 (18.0%) were Butterflies with zero infection, and 126 (14.4%) were Moth with zero infection (Table 1). The distribution of insect species collected per sampling station is presented in Figure 2. The only insect sample infected with EPN (*Mermis* spp.) was from *M. religiosa* during the rainy season. There was no significant difference in EPN infectivity between the nymphs and adult stages in all insect species (P>0.05) across both dry and rainy seasons.

**Figure 2:**
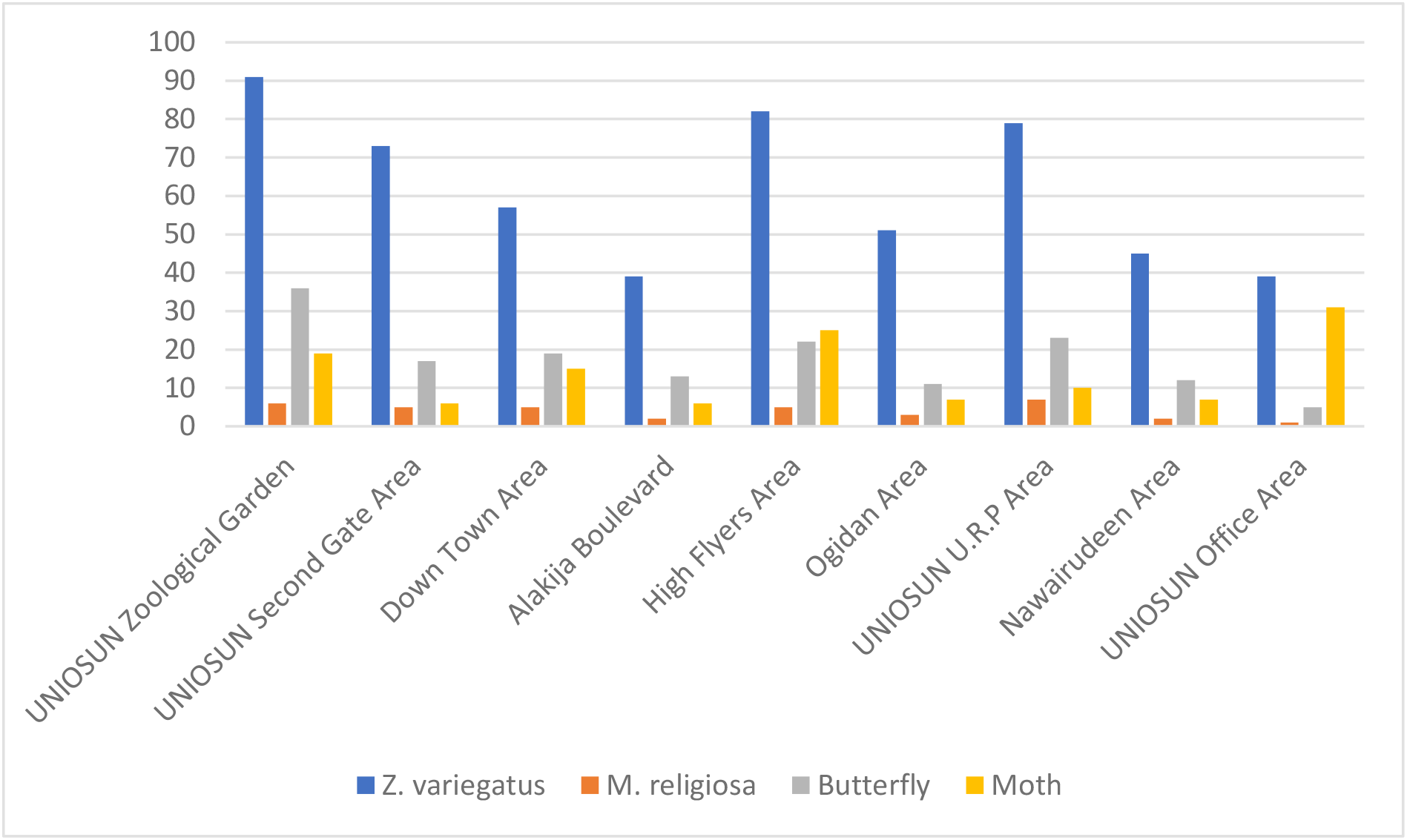
Distribution of sampled insects along sampling stations.

**Table 1:** Prevalence of Entomopathogenic Nematodes (EPNs) in Insect Species Across Sampling Stations in Osogbo Local Govt. Area.

| Species of Insects<br>Collected | Season | Life Stage | No Screened (%) | No Infected (%) | Total Infected<br>(%) |
| --- | --- | --- | --- | --- | --- |
| *Z. variegatus* | Dry | Nymph | 63 (38.2) | 0 | 0 |
|  |  | Adult | 102 (61.8) | 0 |  |
|  | Rain | Nymph | 282 (72.1) | 0 |  |
|  |  | Adult | 109 (27.9) | 0 |  |
|  |  | Total | 556 (63.5) | 0 |  |
| *M. religiosa* | Dry | Nymph | 7 (53.8) | 0 | 1 (0.1) |
|  |  | Adult | 6 (46.2) | 0 |  |
|  | Rain | Nymph | 10 (43.4) | 0 |  |
|  |  | Adult | 13 (56.6) | 1 (7.8) |  |
|  |  | Total | 36 (4.1) | 1(2.8) |  |
| Butterfly | Dry | Larva | 19 (37.3) | 0 | 0 |
|  |  | Adult | 32 (62.7) | 0 |  |
|  | Rain | Larva | 43 (40.2) | 0 |  |
|  |  | Adult | 64 (59.8) | 0 |  |
|  |  | Total | 158 (18.0) | 0 |  |
| Moth | Dry | Nymph | 19 (51.4) | 0 | 0 |
|  |  | Adult | 18 (48.6) | 0 |  |
|  | Rain | Nymph | 31 (34.8) | 0 |  |
|  |  | Adult | 58 (65.2) | 0 |  |
|  |  | Total | 126 (14.4) | 0 |  |
| Overall Total |  |  | 876 |  | 1 (0.1) |

### 3.2 Extraction of Entomopathogenic Nematodes from Soil Samples from Disturbed and Undisturbed Agroecosystems

Out of the 36 soil samples screened for EPNs in the disturbed agroecosystem within Osogbo Local Govt. Area, none of the soil samples were positive for EPN irrespective of their varying soil properties and vegetation (Table 2). However, EPNs are detected in all (24) soil samples collected from undisturbed agroecosystem (Osun-Osogbo Groove) within the same study area with overall *T. molitor larvae* deaths recorded were attributed to EPN infection (Table 3). EPN prevalence in soil samples were not in relation to textural class and vegetation. *Heterorhabditis spp*. dominates all soil samples screened from the undisturbed ecosystem (Table 3). A control was set up for the two sets of experiment, where the *T. molitor* larvae was placed in aired container without soil (Table 2 and 3). The absolute frequency of EPN in soil samples from disturbed agroecosystem was zero whereas 100% absolute frequency was recorded for undisturbed agroecosystem (Table 4).

**Table 2:**
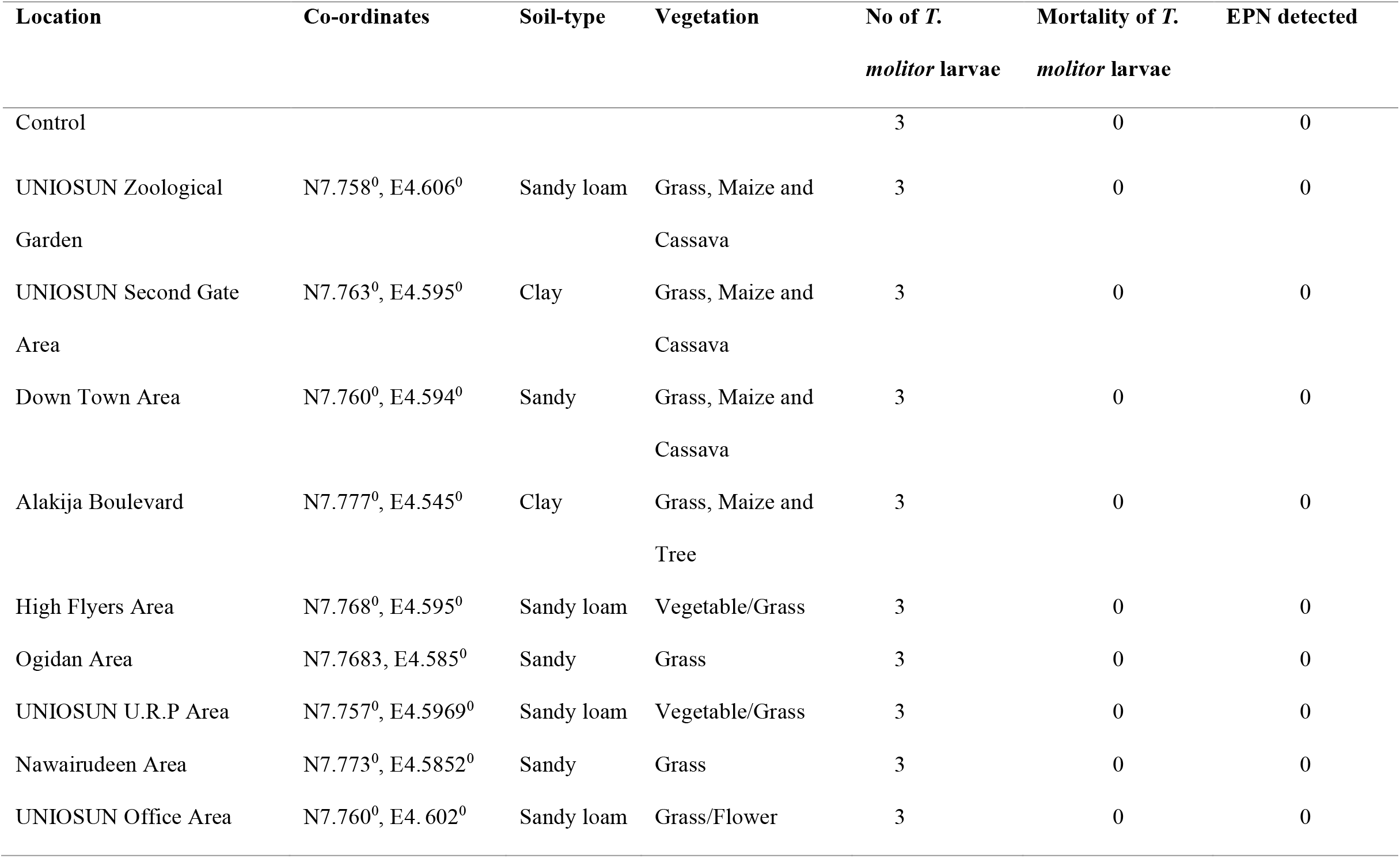
EPN Detection in Soil Samples from Disturbed Agroecosystem Using Baiting Method.

| Location | Co-ordinates | Soil-type | Vegetation | No of *T.*<br>*molitor* larvae | Mortality of *T.*<br>*molitor* larvae | EPN detected |
| --- | --- | --- | --- | --- | --- | --- |
| Control |  |  |  | 3 | 0 | 0 |
| UNIOSUN Zoological Garden | N7.758<sup>0</sup>, E4.606<sup>0</sup> | Sandy loam | Grass, Maize and Cassava | 3 | 0 | 0 |
| UNIOSUN Second Gate Area | N7.763<sup>0</sup>, E4.595<sup>0</sup> | Clay | Grass, Maize and Cassava | 3 | 0 | 0 |
| Down Town Area | N7.760<sup>0</sup>, E4.594<sup>0</sup> | Sandy | Grass, Maize and Cassava | 3 | 0 | 0 |
| Alakija Boulevard | N7.777<sup>0</sup>, E4.545<sup>0</sup> | Clay | Grass, Maize and Tree | 3 | 0 | 0 |
| High Flyers Area | N7.768<sup>0</sup>, E4.595<sup>0</sup> | Sandy loam | Vegetable/Grass | 3 | 0 | 0 |
| Ogidan Area | N7.7683, E4.585<sup>0</sup> | Sandy | Grass | 3 | 0 | 0 |
| UNIOSUN U.R.P Area | N7.757<sup>0</sup>, E4.5969<sup>0</sup> | Sandy loam | Vegetable/Grass | 3 | 0 | 0 |
| Nawairudeen Area | N7.773<sup>0</sup>, E4.5852<sup>0</sup> | Sandy | Grass | 3 | 0 | 0 |
| UNIOSUN Office Area | N7.760<sup>0</sup>, E4.602<sup>0</sup> | Sandy loam | Grass/Flower | 3 | 0 | 0 |

**Table 3:** EPN Verification in Soil Samples from Undisturbed Ecological Setting (Osun-Osogbo Groove) Using Baiting Method.

| Location | Co-ordinates | Soil-type | Vegetation | No of *T. molitor* | Mortality of *T. molitor* | EPN detected |
| --- | --- | --- | --- | --- | --- | --- |
| Control |  |  |  | 3 | 0 | 0 |
| Information Centre | N7°45'29"E4°33'0" | Sandy loam | Trees, Grass | 3 | 2 | *Heterorhabditis* spp. |
| Iyamopo Courtyard | N7°45'15"E4°32'57" | Sandy loam | Trees, Grass | 3 | 2 | *Heterorhabditis* spp. |
| Groove Toilet Area | N7°45'19"E4°33'1" | Sandy loam | Trees, Grass | 3 | 3 | *Heterorhabditis* spp. |
| Mukadan | N7°45'4"E4°33'1" | Sandy loam | Trees, Grass | 3 | 2 | *Heterorhabditis* spp. |
| Artist Village | N7°45'29"E4°32'59" | Sandy loam | Trees, Grass | 3 | 3 | *Heterorhabditis* spp. |
| BMI | N7°45'9"E4°32'59" | Sandy loam | Trees, Grass | 3 | 2 | *Heterorhabditis* spp. |
| Olomoyoyo Courtyard | N7°45'18"E4°33'3" | Sandy loam | Trees, Grass | 3 | 2 | *Heterorhabditis* spp. |
| Elewure | N7°45'10"E4°33'0" | Sandy loam | Trees, Grass | 3 | 3 | *Heterorhabditis* spp. |

\*BMI = A Point Between Mukadan and Iyamopo Courtyard

**Table 4:** Absolute Frequency of EPNs in Both Agroecosystems.

| Location | No. of positive samples | Total samples collected | AF* (%) |
| --- | --- | --- | --- |
| Disturbed Agroecosystem | 0 | 36 | 0 |
| Undisturbed Agroecosystem | 24 | 24 | 100 |

\*Absolute frequency = No. of samples containing entomopathogenic nematode species /Total number of samples collected × 100

## 4. DISCUSSION

This investigation cut across two seasons, dry and rainy seasons. However, the ecological advantage promotes high breeding during the rainy season in the study area. Insect species were abundant in the rainy season compared to the dry season (Table 1). The trend was seen in the abundance of *Z. variegatus*, butterflies, moths, and *M. religiosa* in the rainy season than in the dry season. In both seasons, *Z. variegatus* was the most captured while *M. religiosa* was the least captured insect in the study. This can be associated with the evergreen vegetation of the study area that promotes the breeding of grasshoppers (Riffat, 2018b; Rusconi *et al*., 2016). This study observed that *M. religiosa* exhibited a nocturnal and solitary carnivorous habit, which influenced their low capture frequency. The abundance of the insect species and trap efficiency influenced the overall number of insect species captured in this study. Amongst all insects sampled, only one specimen, a *M. religiosa* species, was infected with an EPN. *Mermis* spp. was identified as the successful EPN species that infected the *M. religiosa* species. The insect was captured while, the EPN species was exiting its body, which gave the first clue for identification. Further identification was made based on its phenotypic characteristics, as described by Baker and Capinera (1997).

Previous studies have reported that the abundance of specific insects boosts the rate of EPN infectivity (Demarta *et al*., 2014; Campos-Herrera, 2015). Ozeum *et al*. (2021) isolated 148 *Mermis* spp. from 226 nematode species, accounting for 65.5% of the total nematodes isolated from 248 insects. However, this study observed a low EPN infectivity in the captured insects. As shown in Table 1, there was no statistically significant difference between the prevalence of EPN species in the sampled insect species between the two seasons (P>0.05). This implies that the abundance of insects and seasonal variation had no impact on the distribution of EPN species in the sampled insects. Further, the study area is characterized by different textural classes, such as sandy loam, clay, and sand (Aizebeokhai *et al*., 2018), with different physico-chemical properties which can influence the occurrence and persistence of EPN species (Bal *et al*., 2017). This study opines, that there was no relationship between textural class of the soil and the occurrence of EPN species in the sampled insects. This could be due to characteristic finer textured of the soil, that exhibited high clay content with smaller pore sizes (Khumalo *et al*., 2021). Similarly, frequent flooding leading to soil erosion may have affected the foraging strategies of EPN species in the study. This phenomenon may have influenced the extremely low infectivity recorded amongst the sampled insects (Kaya & Gaugler, 1993).

The study area is a disturbed agroecosystem which experiences a lot of anthropogenic activities. These anthropogenic activities influence the soil integrity negatively. All farmed lands yielded zero occurrence of EPN which may be due to a myriad of anthropogenic activities bordering on land use/land cover activities. However, this observation contradicts the work of Rufai *et al*. (2020) who reported high occurrence of EPN in the study area. This ecological disturbance can be linked to the high anthropogenic activities in the study area (Campos-Herrera *et al*., 2012). Conventional tillage regime, inorganic fertilizer application, crop rotation, and variety selection, bush-burning, to mention a few, are practiced in the study. These management practices are known to considerably affects the diversity of EPN species and their relationship with other organisms (Campos-Herrera *et al*., 2012), which envisage the reason for their disappearance.

It was observed that the effect of the conventional tillage regime practiced in the study area exceeds beyond disrupting the physico-chemical state of the soil but also its biological properties (Neher *et al*., 2019). This is in agreement with the work of Briones and Schmidt (2017), who reported that the conventional tillage regime led to the drop in soil faunal biomass. Also, this is in line with the report of Campos-Herrera *et al*. (2008), who reported that EPNs are less detected in conventional tillage regimes but more in reduced tillage. One of the adverse effects of recurrent bush-burning practices on soil community is the significant interference with the biodiversity in the ecosystem, which is another important cue to the loss of EPN species in the study (Otitoju *et al*., 2019). More so, crop rotation is highly practiced in the study area. However, this could be another cause of EPN suppression, as studies have shown that crop rotation and variety practices influence the food web complexity and consequently has a significant impact on the nematode community in the soil (Neher *et al*., 2019). Neher *et al*. (2019) confirmed that crop rotation could lead to the natural suppression of certain nematode species while others are favoured. Furthermore, the application of inorganic fertilizer to boost crop yields is often practiced. Previous studies have indicated that inorganic fertilizers affect EPN occurrence by reducing their infectivity and virulence in the soil (Jaffuel *et al*., 2016, 2017). The lifespan of crops and varieties cultivated in the study area, which are commonly bi-annual and annual crops, is suggested to have also, impacted the activities of EPN, as earlier research has reported that EPN activity tends to be poor in annual crops (Campos-Herrera *et al*., 2008, 2015). The interaction of these management practices coupled with the aforementioned soil parameters greatly influence the natural occurrence and persistence of EPNs in any geographical location (Alumai *et al*., 2006).

Further verification was made to confirm the claims stated above by this current investigation—soil samples from an undisturbed ecosystem which falls within the territory of the Osogbo Local Government Area; Osun-Osogbo Sacred Groove was screened for EPN (Table 3). This ecosystem was in its natural state, with fewer anthropogenic activities. The results confirmed the presence of EPN species and high as 100% absolute frequency (Table 3 and 4). Of all the EPN isolates recovered, *Heterorhabditis* spp. was the major EPN species identified from the soil samples. Although, EPN species are known to occur and persist in relation to soil type and habitat, especially in sandy loam soils, which aids their mobility and survival, this current prevalence can be attributed to fewer anthropogenic activities, resulting in their spatial distribution within this site (Rufai *et al*., 2020). This proved that EPN species’ disappearance from the disturbed agroecosystem results from intense anthropogenic activities over time. This claim will not be out of place since both the disturbed and undisturbed ecosystem shared close characteristics such as seasonality, vegetation cover, textural class, and the same geographical area (Table 2 and 3).

The result from this study is in line with previous study done by Bal *et al*. (2017). The work of Bal *et al*. (2017) have proposed that certain soil management practices influenced the spatial distribution of EPN infective juveniles in the soil over a period of two years. The results of this study show that anthropogenic activities such as conventional tillage regimes, bush-burning, and inorganic fertilizer application may help predict nematode occurrence. Thus, EPNs are likely to be found in sites with fewer management activities. Similarly, the consequences of global environmental changes, known to influence the biology and ecology of soil biota (García-Palacios *et al*., 2015), cannot be left out of this discussion. Liu *et al*. (2019) indicated that how these global changes influence the spatial distribution of nematode populations, diversity, and functional traits remains obscure. He argued that many climatic events occur in various ways throughout the globe, and in light of the spatial heterogeneity of disturbance, soil biota may end up in a range of more or less suitable environments.

Nonetheless, this study has shown that understanding EPN’s ecology is vital for predicting their suitability against target insect species. With the changing dynamics of EPN species due to anthropogenic activities and other natural events, it is crucial to carry out further studies using molecular ecology to provide insights into EPN dispersion, variation, and genetic adaptation, which would form a build-up knowledge for exploring other essential traits related to their survival and infectivity beyond soil parameters and foraging strategies. Also, further studies into the effects of biotic factors on EPNs survival and reproduction are needed to be explored. All these are necessary to be considered as they would boost the efficiency of these beneficial organisms against target insect pests.

## 5. CONCLUSION

EPN behavioural and survival strategies are more complicated beyond their foraging strategies and level of infectivity on target organisms. Therefore, this study states that intense anthropogenic activities adversely affect EPN occurrence, dispersal, and persistence in the soil. These findings present EPN as a potential tool for environmental impact assessment, as their presence and disappearance can be linked to specific environmental parameters and human activities. However, this study backs the standing hypothesis that soil management practices would influence the spatial distribution of EPN infective juveniles in the soil over two years. The disappearance of EPN species from an area known to persist is a call for environmental impact assessment. This study, therefore, infers and concludes that Osogbo Local Govt. Area of Osun State is an area facing serious ecological disturbance arising greatly from unchecked human activities.

